# Efficient mutagenesis of maize inbreds using biolistics, multiplex CRISPR/Cas9 editing, and Indel-Selective PCR

**DOI:** 10.1101/2024.10.21.619474

**Authors:** Maruti Nandan Rai, Brian Rhodes, Stephen Jinga, Praveena Kanchupati, Edward Ross, Shawn R. Carlson, Stephen P. Moose

## Abstract

CRISPR/Cas9 based genome editing has advanced our understanding of a myriad of important biological phenomena. Important challenges to multiplex genome editing in maize include assembly of large complex DNA constructs, few genotypes with efficient transformation systems, and costly/labor-intensive genotyping methods. Here we present a ‘fast-edit’ approach for multiplex CRISPR/Cas9 genome editing system that delivers a single compact DNA construct *via* biolistics to type I embryogenic calli, followed by a novel efficient genotyping assay to identify desirable editing outcomes. We first demonstrate the creation of heritable mutations at multiple target sites within the same gene. Next, we successfully created individual and stacked mutations for multiple members of a gene family. Genome sequencing found off-target mutations are rare. Multiplex genome editing was achieved for both the highly transformable inbred line H99 and Illinois Low Protein1 (ILP1), a genotype where transformation has not previously been reported. In addition to screening transformation events for deletion alleles by PCR, we also designed PCR assays that selectively amplify deletion or insertion of a single nucleotide, the most common outcome from DNA repair of CRISPR/Cas9 breaks by non-homologous end-joining. The Indel- Selective PCR (IS-PCR) method enabled rapid tracking of multiple edited alleles in progeny populations. The ‘end to end’ pipeline presented here for multiplexed CRISPR/Cas9 mutagenesis can be applied to accelerate maize functional genomics in a broader diversity of genetic backgrounds.

## Introduction

Maize is the global leader for grain production, with more than one billion tons harvested annually (Erenstein *et al*., 2022). Evaluation of natural variation in industrial breeding programs has led to dramatic improvements in maize yields worldwide (Wang *et al*., 2020). The recent burst of genome data and functional genomics resources for maize has uncovered multiple novel targets for further improvement. CRISPR/Cas9 technology is utilized to generate mutations in targeted genomic regions in many plant species including maize (Varotto, 2024). However, multiplexing CRISPR/Cas9 genome editing presents a number of technical challenges, including assembly of constructs with multiple DNA elements, a limited number of RNAP III promoters typically used to drive sgRNA expression, inefficient transformation protocols, and subsequent genotyping to identify and follow inheritance of edited mutations.

One solution to reduce size and complexity of editing constructs is to express multiple sgRNA scaffolds as one polycistronic mRNA separated by processing elements, each driven by a single RNA polymerase II promoter. In addition to simplifying construct assembly using Type IIS restriction-mediated cloning approaches such as GoldenGate, RNA polymerase II promoters allow for targeted expression in certain tissues or at precise developmental times (Tang *et al*., 2016) and are thus more versatile than RNA polymerase III promoters. In one example of this approach, Qi et al., (2016) utilized the existing tRNA processing pathway in plants and introduced flanking tRNA processing sites within a transcript containing multiple guide RNAs driven by a single RNA polymerase II promoter. Two additional multiplexing strategies that have been shown to function in plants are self-cleaving ribozymes (He *et al*., 2017) and the Csy4 ribonuclease that mediates bacterial processing of CRISPR arrays (Nissim et al., 2014; McCarty et al., 2020; Hassan et al., 2021). A compact vector system has been developed for assembly and Csy4 processing of multiple gRNAs expressed from the constitutively active Cestrum yellow leaf curling virus promoter (Čermák et al., 2017), with successful demonstration of multiplexed editing in crops such as soybean (Luo *et al*., 2021), rapeseed (Wang *et al*., 2021), and tomato (Shiose *et al*., 2024).

Another bottleneck in CRISPR/Cas9 mutagenesis of maize is the requirement to deliver editing components to regenerable tissue cultures, which can be reliably induced from only a few genotypes. The most efficient maize genotype for *Agrobacterium*-mediated transformation, Hi-II, is a hybrid, potentially complicating phenotypic analysis of progeny. First reported by Lowe et al., (2016, 2018) and with recent improvements (Wang *et al*., 2023), use of the morphogenic regulators BABYBOOM and WUSCHEL dramatically expands the genotypes available for transformation. However, these methods still require careful optimization of expression, with some of the key elements being governed by intellectual property that may limit access by academic groups.

Even when multiplex genome editing is successful, efficient identification of individual alleles among the initial population of mutations or their inheritance among progeny can be challenging. Although genotyping ‘drop-out’ mutations (deletion of >10bp) can be done easily by simple PCR and gel electrophoresis, such assays do not reveal single nucleotide insertion/deletion (indel) mutations that are the most common editing outcome in plants (Bortesi *et al*., 2016). Both DNA sequencing and *in vitro* RNP CRISPR/Cas9 digestion assays of amplicons spanning gRNA target(s) can readily identify single-base edits, but are not cost-effective for large-scale screening or introgression of CRISPR/Cas9 derived alleles into additional genetic backgrounds of interest.

The current work presents an approach aimed to address each of the above challenges for multiplex genome editing in maize. Multiple gRNAs were cloned into the compact Csy4 vector system developed by Čermák *et al*. (2017). Single DNA fragments containing each of Cas9, gRNA scaffolds, and selectable marker gene were delivered via biolistics to Type I embryogenic calli, which can be established from a much broader range of maize genotypes (Duncan et al., 1985). We demonstrate successful multiplexed genome editing in two different maize inbred lines by creating heritable mutations at multiple target sites within the same gene, as well as multiple members of a multigene family. Lastly, we developed a novel and efficient PCR-based assay to selectively amplify single-base indel mutations, and apply Indel-Selective PCR to genotype populations for desirable combinations of edited alleles.

## Results

### DNA design for multiplex editing of the maize *Lemon White1* gene

Multiplex editing is achieved with either multiple constructs expressing individual components (nuclease, gRNAs, selectable marker) or single constructs that express all components. Constructs expressing individual components are easier to build and offer more flexibility in design, but the frequency of codelivery decreases with the number of constructs. Single constructs are desired but can present difficulties in the cloning and propagation of large (often greater than 15-kb) inserts with repeated elements, such as RNA polymerase III promoters or gRNA scaffolds. The vectors designed by Čermák *et al*. (2017) address many of these issues by expression of multiple gRNAs as a polycistronic mRNA, driven by the compact CMYLV promoter (Čermák *et al*., 2017). Expression of an N-terminal fusion of the Csy4 processing protein to a monocot codon-optimized Cas9, separated by the P2A self-cleaving peptide, is controlled by the strong constitutive *ZmUbi1* promoter. The editing components are coupled to a selectable marker gene for recovery of transgenic events. GoldenGate entry sites facilitate both direct cloning of multiple synthesized gRNA scaffolds into a common backbone, or modular assembly of different promoters and editing components.

To assess functionality of the Csy4 multiplexed editing system in maize, we selected the *Lemon White 1 (LW1,* Zm00001eb056240) gene for a proof-of-concept experiment because of prior success in generating easily visualized yellow/albino leaf phenotypes by CRISPR/Cas mutagenesis (Feng et al., 2016; Trieu et al., 2022). Four guide RNAs were designed to target *LW1* exons using the B73 reference genome assembly (Supplementary Table 1). LW1-gRNA1 generated edits previously (Feng *et al*., 2016) was included as a positive control (Figure 1A), along with three gRNAs targeting either upstream (LW1-gRNA2, LW1-gRNA4) or downstream (LW1- gRNA3) exons (Figure 1A). All four LW1-gRNAs were assembled into the pMOD_B2103 destination vector, which was then combined with plasmids containing ZmUbi1:Csy4-TaCas9 and the CaMV35S-*nptII* selectable marker (Figure 1B). The editing function of this vector was initially tested with *in vitro* Cas9 RNP cleavage assays, (supplemental Figure 1) where each of the three newly designed gRNAs produced expected cleavage products when tested individually, as well as in combination.

**Figure 1:**
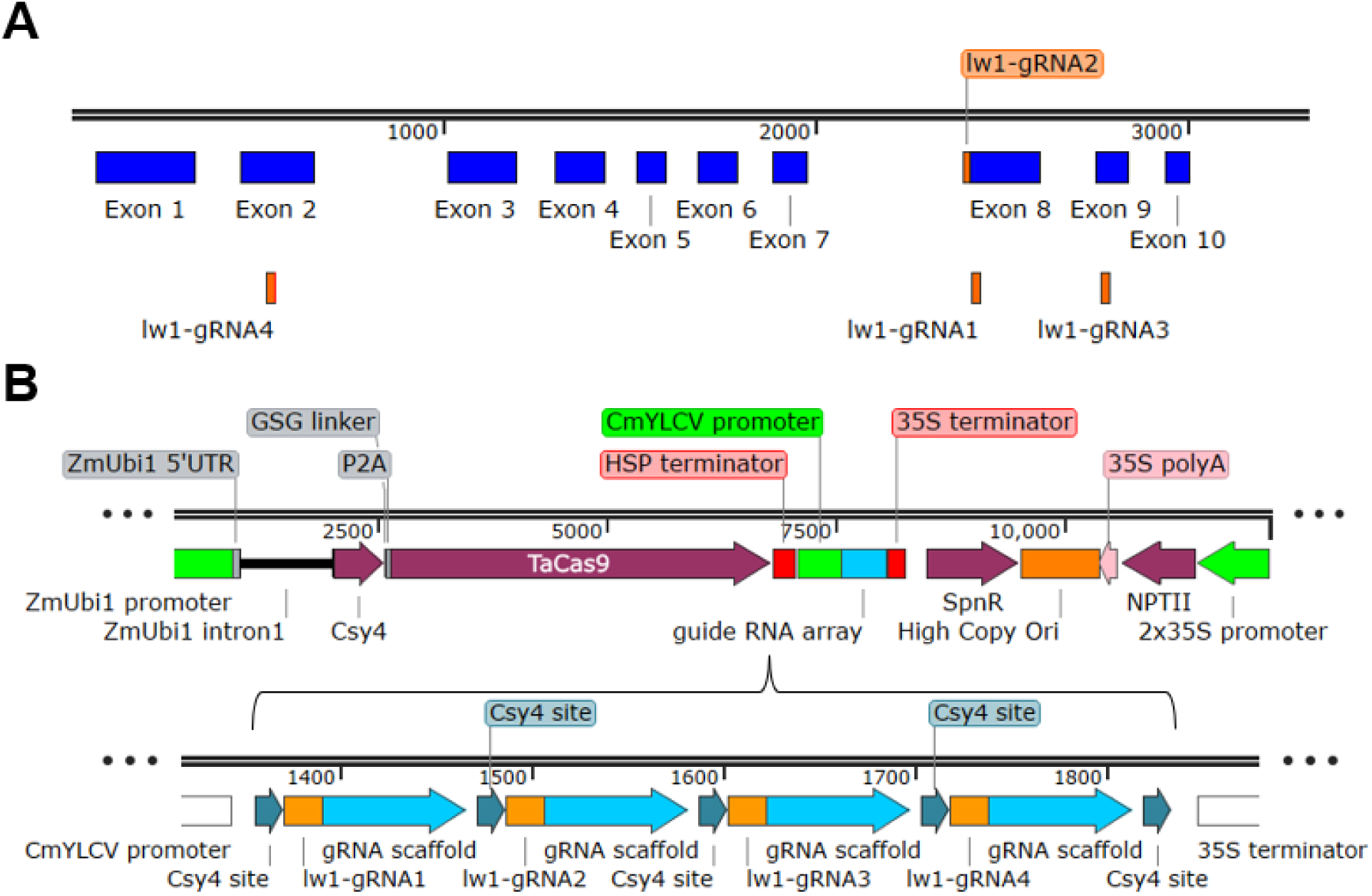
DNA design for multiplex editing of the maize *Lemon White1* gene. **(A)** *Lemon White 1* (*LW1*) gene model depicting exons and the position of designed guide RNAs. **(B)** Schematic design of single Editing DNA construct delivered by biolistics for multiplex *LW1* gene editing.

### Multiplex editing of the *LW1* gene in the H99 and ILP1 maize inbred lines

The two most common methods for DNA delivery in maize transformation are biolistics or *Agrobacterium*. The first fertile transgenic maize plants were produced using biolistics (Gordon-Kamm *et al*., 1990). Once protocols were developed for *Agrobacterium* transformation of maize (Ishida *et al*., 1996), it became favored for DNA delivery due to the higher frequency of single-copy transgenic events. However, efficient *Agrobacterium* transformation remains limited to only a few maize genotypes. When generating mutations by CRISPR/Cas9, fewer independent events are required compared to transgene expression and transgenes will eventually be removed by genetic segregation. Furthermore, the ability to deliver higher concentrations of DNA encoding editing components on microprojectiles compared to *Agrobacterium* may increase editing efficiency. Biolistic methods were used to create mutations at the *waxy* locus in a number of elite maize inbred lines (Gao *et al*., 2020) demonstrating the versatility of this approach. DNA delivery by biolistics opens the possibility to apply editing to the broader diversity of maize inbred lines that are capable of forming embryogenic callus. Accordingly, we sought to develop an editing pipeline using biolistics delivery to maize immature embryos or Type I callus that was subsequently regenerated into plants.

Among maize inbred lines, H99 has historically been transformed at higher efficiencies compared to other inbred lines when using biolistics delivery (Brettschneider et al., 1997; Shiva Prakash et al., 2008). H99 is classified as a Non-Stiff Stalk inbred and thus can form high-yielding hybrids when combined with Stiff Stalk parents such as B104 that is a preferred genotype for maize editing (Aesaert *et al*., 2022; Kang *et al*., 2022). To demonstrate the versatility of the biolistics- Type I callus transformation system, we also chose to generate mutations in the inbred line Illinois Low Protein1 (ILP1). ILP1 is derived from the Illinois Long-Term Selection Experiment for grain protein concentration (Moose et al., 2004), exhibits high nitrogen utilization efficiency (Uribelarrea *et al*., 2007) and harbors novel genetic variants for improving maize yield, seed composition, and nitrogen use efficiency (Zhang *et al*., 2019). Previous work showed that Type I embryogenic callus and successful plant regeneration can be achieved with the source population for ILP1 (Duncan *et al*., 1985).

The single 12-kbp DNA fragment for *Lw1* editing (Figure 1B) was used to transform embryogenic calli of the H99 and ILP1 genotypes by biolistic transformation. Following methods initially described in and further optimized by Shiva Prakash *et al*.. (2008), transgenic events harboring *lw1* mutations were generated for both H99 and ILP1 (Figure 2). As anticipated, embryogenic calli induced from ILP1 exhibited the morphogenic features of Type I embryogenic calli (Figure 2B). Approximately 12 weeks post-bombardment, putative transgenic shoots appeared on selective regeneration medium (Figures 2C). Table 1 shows transformation efficiency of H99 in this experiment was comparable to prior reports (4.5%), and a slightly higher frequency of 7.2% was observed for ILP1. Importantly, more than 20 events were produced for both genotypes. Transgenic plantlets were observed to display either partial or fully albino leaves (Figure 2) indicative of *lw1* mutations.

**Figure 2.**
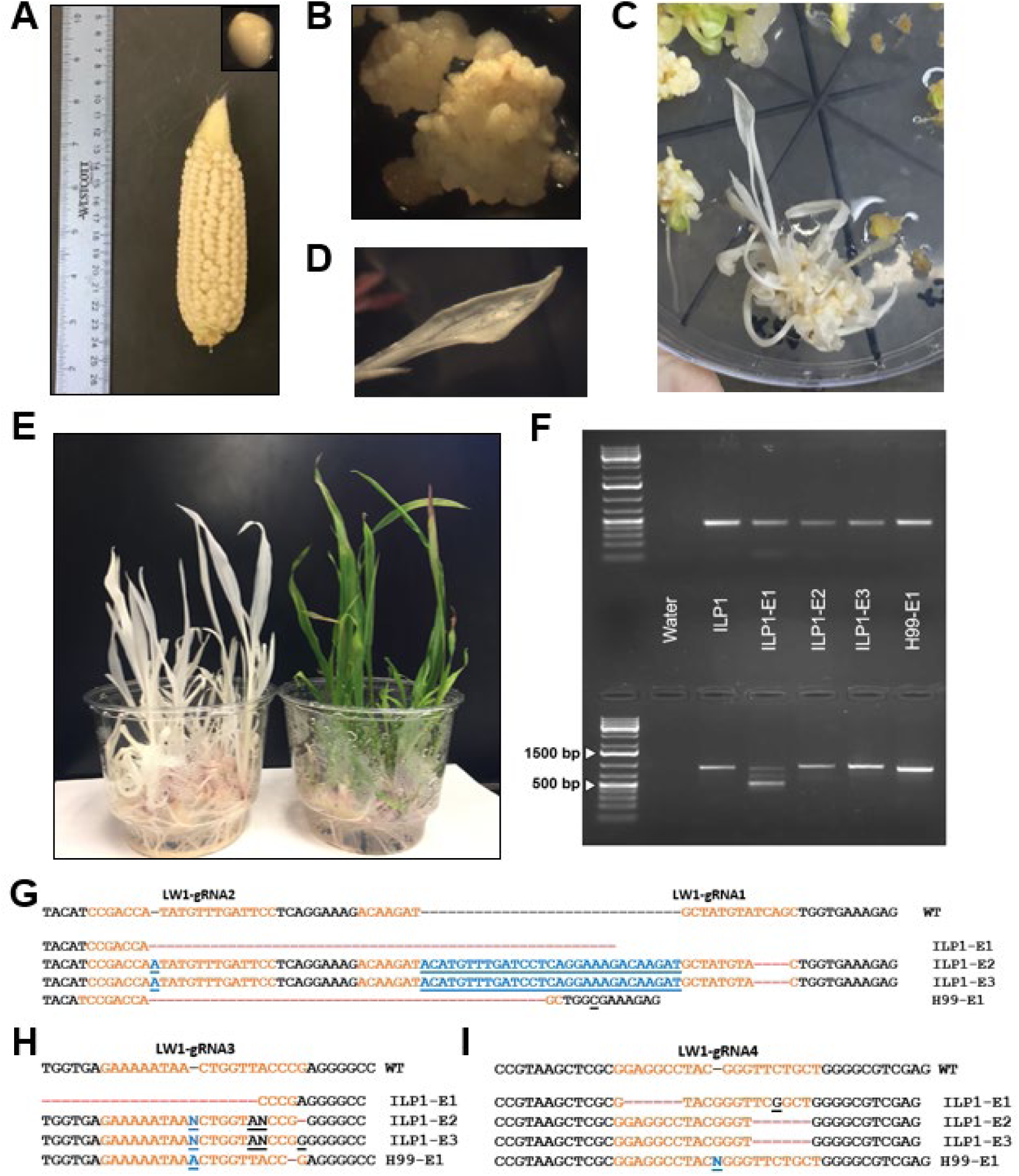
Multiplex genome editing of *LW1* in H99 and ILP1 inbred lines. **(A)** Selfed ears of ILP1 harvested 12-14 days after pollination, with excised immature embryo in inset. **(B)** Putative transgenic calli on selective medium**. (C)** CRISPR/Cas9-events on selective regeneration medium. **(D, E)** Putative transgenic shoots on selective regeneration medium after 2- 3 weeks post-transfer to a growth chamber with 16h/8h light/dark photoperiod. **(F)** Sequence confirmation of genome edits in *LW1*. The top half of the gel shows amplification of the 5’ end of the gene spanning the *LW1*-gRNA4 target site, bottom half shows amplicons spanning target sites for *LW1*-gRNA1, *LW1*-gRNA2 and *LW1*-gRNA3 at 3’ end of gene. Left lane in each gel is the NEB 1kb+ ladder. **(G, H, I)** Sanger sequencing results of PCR products shown in panel F (guide RNA targets are depicted in orange, PAM sequences are depicted in bold black letters; red dashes represent deletions, bold blue underlined letters represent insertions, and base changes are depicted in black underlined letters).

**Table 1.**
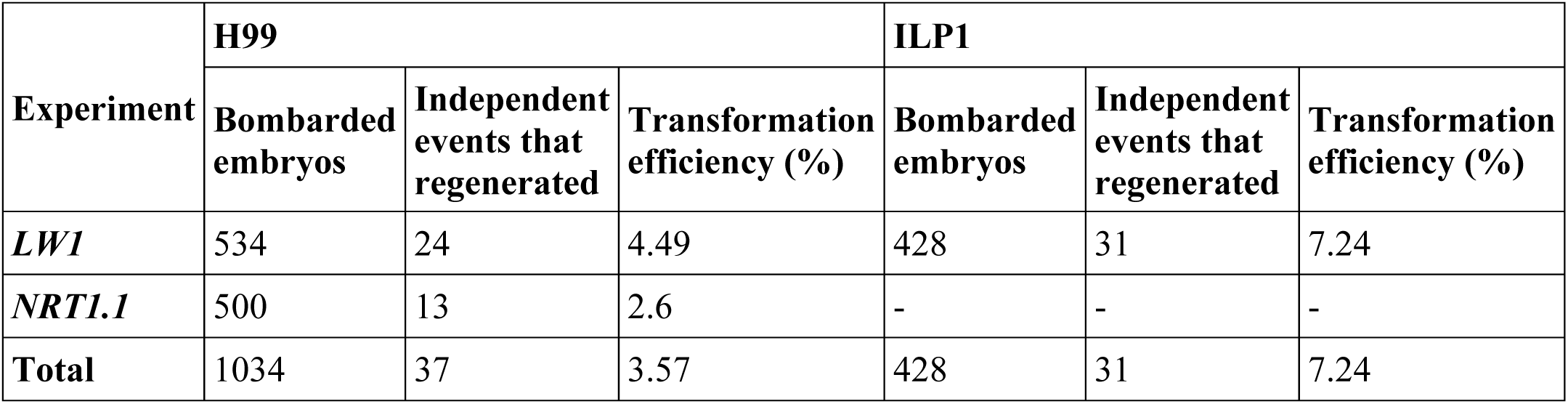
Transformation efficiency of independent experiments with immature embryo- derived calli of H99 and ILP.

PCR amplification of genomic DNA isolated from putative *lw1* mutant tissue followed by gel electrophoresis revealed successful multiplex editing, by producing deletion mutations at two *LW1* target sites. Data from three events are presented as examples. ILP1 event 1 (ILP-E1) produced an amplicon ∼500 bp in length, a size consistent with the expected loss of 385 bp between *LW1*-gRNA2 and *LW1*-gRNA3 (Figure 2F). Sanger sequencing of PCR amplicons that are the same size as the untransformed control further confirmed mutations at each of the genomic sequences targeted by gRNAs (Figure 2G-I). Two separate plants regenerated from ILP1 event 2 (ILP-E2) contained the same mutations at each of the four *LW1*-gRNA target sites. For one event in the H99 background (H99-E1), an amplicon of ∼800 bp was produced because of a 29-bp deletion generated following cleavage at the *LW1*-gRNA1 and *LW1*-gRNA2 sites (Figure 2F &2G). To summarize, Csy4 multiplexed CRISPR/Cas9 editing created multiple distinct mutations in all four gRNA target sites in the *LW1* gene.

### Mutagenesis of a gene family

The experiments with *LW1* demonstrated each of the construct elements was functional in multiplex genome editing. We next examined if we could use this method to ‘fast-edit’ multiple genes of a family using guide RNAs designed to target sequences conserved across all members, or unique to individual genes. Because of our interest in nitrogen utilization, we decided to edit the *NRT1.1* gene family of nitrate transporters (Figure 3A). Maize contains four *NRT1.1* family members (Plett *et al*., 2010). *NRT1.1A* (Zm00001d024587) is located on chromosome 10, *NRT1.1B* (Zm00001d029932) and *NRT1.1C* (Zm00001d029933) are tandemly-duplicated copies on chromosome 1, and *NRT1.1D* (Zm00001d027285) resides at the distal tip of the short arm of chromosome 1. Inspection of gene expression data for these genes at MaizeGDB.org shows that compared to *NRT1.1A* and *NRT1.1B*, *NRT1.1C* is weakly expressed and no expression is observed for *NRT1.1D*, which is likely a pseudogene. These expression differences were also observed for leaf tissue from multiple maize genotypes (Cheng *et al*., 2021). To simultaneously generate edits in each of the *NRT1.1A*, *NRT1.1B* and *NRT1.1C* genes, we designed three gRNAs: NRT1.1- gRNA2 targeted a perfectly conserved sequence in each gene, whereas NRT1.1-gRNA1 and NRT1.1-gRNA3 specifically targeted *NRT1.1B* (Figure 3A). To design NRT1.1-gRNA2, the off-target requirement was relaxed to allow perfect matching at the three genes. All guide RNA targets are located within or upstream of several transmembrane domains within NRT1.1 proteins that are core for small molecule transport. The functionality of the NRT1.1-gRNAs at each of the three NRT1.1 gene targets was verified using both *in vitro* Cas9 RNP assays (data not shown) and after transformation of maize leaf protoplasts (Supplementary Figure 2).

**Figure 3:**
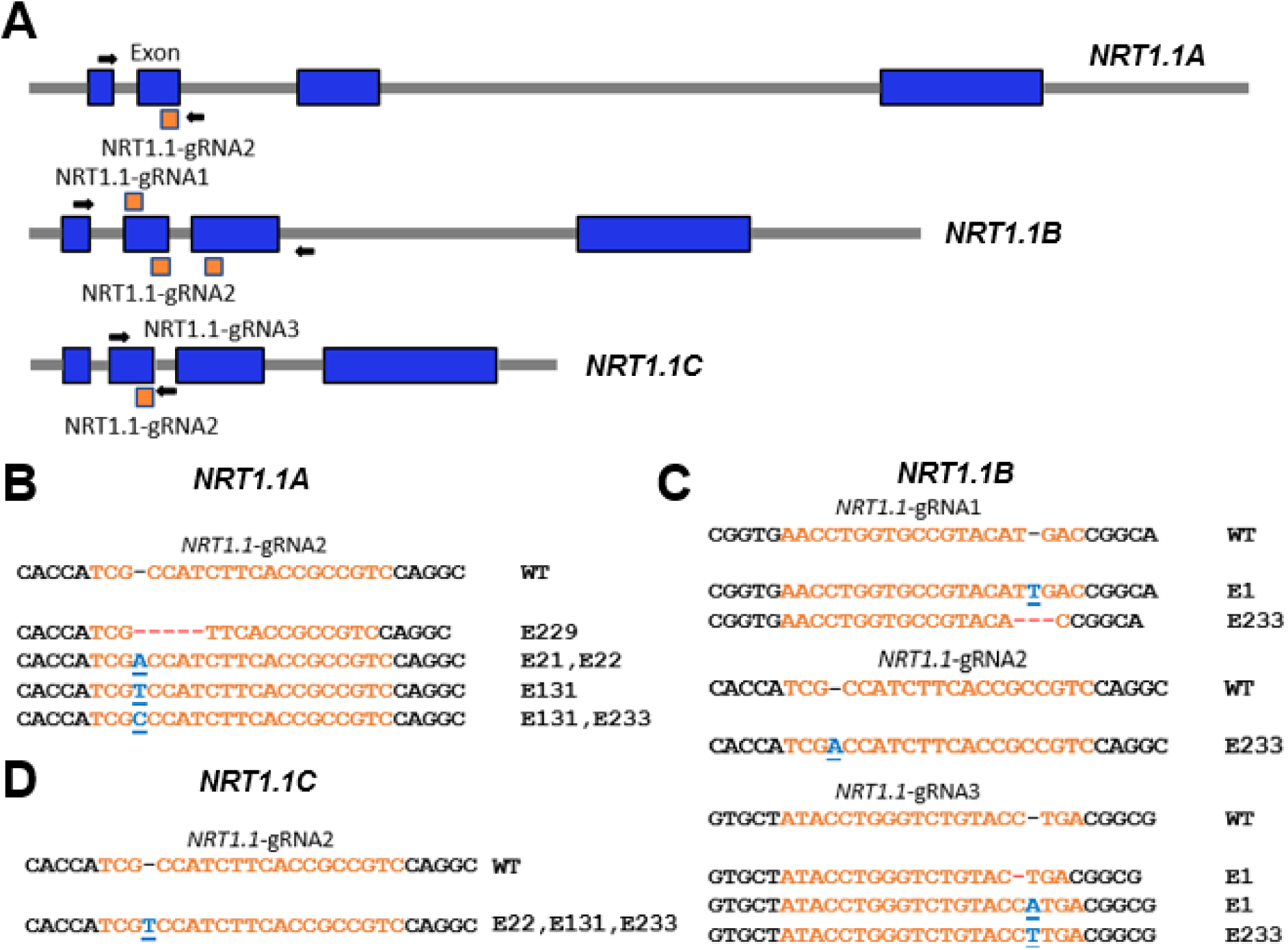
Mutagenesis of the maize *NRT1.1* gene family by multiplex CRISPR/Cas9 editing. **(A)** *NRT1.1 A*, *B* and *C* gene models depicting exons (blue boxes), introns (grey lines) and the target sites for designed guide RNAs (orange boxes). The arrows on the gene models represent the gene-specific primers used for PCR confirmations of edits in plants**. (B, C & D)** Sanger sequencing results of PCR products amplified using gene specific primers in gene-edited plants (guide RNA targets are depicted in orange, red dashes represent deletions, bold blue underlined letters represent insertions).

Immature H99 maize embryos were bombarded with the NRT1.1 multiplex editing construct and thirteen resistant calli were recovered, resulting in a transformation efficiency of 2.6% (Table 1). Of these 13 events, seven were confirmed to contain a mutation at one or more NRT1.1-gRNA target sites. All seven events were successfully regenerated into fertile T0 plants. Table 2 summarizes the spectrum of mutations obtained in the *NRT1.1* gene family that were identified by Sanger sequencing of PCR amplicons from the seven independent callus lines and three regenerated T0 plants from each event (Figure 2B, 2C & 2D, Table 2). Overall, 19 different mutations were recovered from the three *NRT1.1* genes (Table 2). Among these mutations, 12 were an insertion of one or two base pairs and two others were small deletions of 1-18 base pairs. T0 edited plants ranged from chimeric, heterozygous, or homozygous for bi-allelic mutations.

**Table 2:** DNA sequence changes in callus and subsequently regenerated plants for *NRT1.1A*, *NRT1.1B* and *NRT1.1C* genes within each event confirmed to contain the Cas9 protein, guide RNA array and selection marker gene. Mutations occurring at the locations of the three gRNAs targeting *NRT1.1B* are listed in order of *NRT1.1*-gRNA1, *NRT1.1*-gRNA2 and *NRT1.1*-gRNA3. Estimated proportion of edited DNA (%) in callus (according to TIDE analysis) is also listed. (Abbreviation - WT: wild type; Hetro- heterozygous for the mutation; UE: unknown edit).

| *NRT1.1A* |  |  |  |  |  |
| --- | --- | --- | --- | --- | --- |
| Events | mutation type | Callus<br>estimated proportion<br>of edited DNA (%) | Plants |  |  |
|  |  |  | 1 | 2 | 3 |
| 1 | +1 (C) | 46.7 | WT | WT | WT |
| 21 | +1 (A) | 46.5 | Het +1 (A) | Het +1 (A) | Het +1 (A) |
| 22 | +1(A),<br>+1(T) | 50, 50 | Het +1 (A) | Het +1 (A) | Het +1 (A) |
| 70 | WT | 100 | WT | WT | WT |
| 131 | +2 | 16.2 | Het +1 (C) | WT | Het +1 (T) |
| 171 | +1 (A) | 10.1 | N/A | N/A | N/A |
| 229 | -5 | 45.1 | Het -5 | Het -5 | Het -5 |
| 233 | +1 (C) | 100 | +1 (C) | +1 (C) | +1 (C) |

| *NRT1.1B* |  |  |  |  |  |
| --- | --- | --- | --- | --- | --- |
| Events | mutation type | Callus<br>estimated proportion<br>of edited DNA (%) | Plants |  |  |
|  |  |  | 1 | 2 | 3 |
| 1 | WT | 0 | UE | UE | WT |
| 21 | WT | 0 | WT | WT | WT |
| 22 | UE | N/A | WT | WT | WT |
| 70 | WT | 0 | WT | WT | WT |
| 131 | UE | N/A | UE | WT | UE |
| 171 | WT | 0 | N/A | N/A | N/A |
| 229 | WT | 0 | WT | WT | WT |
| 233 | UE | N/A | -3, +1(A),<br>+1(T) | -3, +1(A),<br>+1(T) | -3, +1(A),<br>+1(T) |

| *NRT1.1C* |  |  |  |  |  |
| --- | --- | --- | --- | --- | --- |
| Events | mutation type | Callus<br>estimated proportion<br>of edited DNA (%) | Plants |  |  |
|  |  |  | 1 | 2 | 3 |
| 1 | -18, +2 | 51.1, 45.3 | WT | WT | WT |
| 21 | +1 (C) | 54.8 | WT | WT | WT |
| 22 | +1(A),<br>+1(T) | 50, 50 | Het +1 (T)* | Het +1(T) * | WT |
| 70 | WT | 0 | WT | WT | WT |
| 131 | +1 (T) | 21.9 | Het +1 (T) | WT | Het +1 (T) |
| 171 | +1 (T) | 8.4 | N/A | N/A | N/A |
| 229 | WT | 0 | WT | WT | WT |
| 233 | +1 (T) | 100 | +1 (T) | +1 (T) | +1 (T) |

Eight different *nrt1.1a* mutations were generated, with a single ‘C’ insertion occurring in two events (E1 and E233) and a single ‘A’ insertion in both events 21 and 171 (Table 2). Three events (E22, E131 and E233) showed evidence of editing at *NRT1.1B*, but we were not able to precisely identify the nucleotide differences (Table 2). Eight different mutations were observed in *NRT1.1C*, including a single ‘T’ insertion found in four events (E22, E131, E171, and E233; Table 2). Among the 19 mutations observed in calli, 9 were also observed in DNA from seedling leaves of the regenerated T0 plants, at least three mutations for each of *NRT1.1A*, *NRT1.1B* and *NRT1.1C* (Table 2, Figure 3B, 3C & 3D). Conversely, two new single base insertions were identified in T0 plants for *NRT1.1A* (E131; Figure 3B) and another in *NRT1.1B* near *NRT1.1*-gRNA3 (E1; Figure 3C) that were not identified at the callus stage (Table 2). At the T0 plant stage, the “unknown” edits in *NRT1.1B* for the callus event 233 was resolved to a homozygous mutation comprised of a 3-bp deletion near *NRT1.1*-gRNA1, a single ‘A’ insertion near *NRT1.1*-gRNA2 and a single ‘T’ insertion at *NRT1.1-*gRNA3 (Table 2; Figure 3C). The single base insertions in Event 21 (for *NRT1.1A*) and Event 131 (for *NRT1.1C*), as well as the 5-bp deletion for *NRT1.1A* found in event 229, each showed approximately 50% editing in both callus and T0 plants, indicative of fully heterozygous edits (Table 2; Figure 3B, 3C & 3D). Importantly, event 233 showed homozygous edits at all possible guide RNA target sites in callus and all 3 T0 plants, indicating Csy4 based multiplex editing of the *NRT1.1* gene family could proceed to completion in a single transformation event (Figures 3B, 3C & 3D).

The heritability of the *NRT1.1* gene family edits was assessed in T1 progeny produced from either self-pollination of T0 plants or crossing as either male or females with control H99 plants. The inheritance of edited alleles in T1 progeny was confirmed by Sanger sequencing of the PCR amplicons using primers flanking identified mutations in T0 plants from events E1 and E233 (Figure 3B-D). All together these results demonstrate that multiplexed Csy4 CRISPR/Cas9 editing system successfully created mutations in multiple genes of a multi-gene family.

### Genome sequencing reveals rare off-target mutations in maize genome

To identify potential off-target mutations generated by the *NRT1.1* genome editing system, we performed Illumina deep sequencing on T0 seedling leaf tissue from event 1. This sequencing run produced 70 million total paired reads which gives an average read depth of 8.75X across the maize genome. These sequence reads were aligned to Oh43, which is the most closely related inbred to H99 with a complete genome assembly (Hufford *et al*., 2021). Each of the 20 genomic regions predicted as off target sites for the three NRT1.1-gRNAs were visually inspected for sequence variation indicative of editing. The average read depth within these off-target regions was 13.6. When allowing up to three sequence mismatches, only one showed evidence of off-target mutagenesis (Table 3), which coincided with *NRT1.1D*, the 4^th^ member of the *NRT1.1* gene family (Figure S3A). This putative edit contained a one base-pair mismatch 17 nucleotides from the PAM proximal region of the NRT1.1-gRNA2 target sequence (Figure S3B). The read depth at this site was 12 and 4 of these reads showed an A nucleotide insertion, resulting in a 33.3% editing rate. In conclusion, off-target analysis indicated a rare occurrence of off target mutagenesis, and only at the locus already known to share the strongest match with intended editing targets.

**Table 3:** Frequency of off-target editing for the NRT1.1 gene family. Potential off-target sequences containing either 1, 2 or 3 mismatches to the three *NRT1.1* guide RNAs and number showing evidence of editing based on Illumina deep sequencing.

|  | gRNA1 | gRNA2 | gRNA3 |
| --- | --- | --- | --- |
| **Off-targets** | 1 mismatch |  |  |
| Possible: | 0 | 3 | 0 |
| Edited: | 0 | 1 (33% of all reads) | 0 |
|  | 2 mismatches |  |  |
| Possible: | 2 | 2 | 2 |
| Edited: | 0 | 0 | 0 |
|  | 3 mismatches |  |  |
| Possible: | 4 | 5 | 2 |
| Edited: | 0 | 0 | 0 |

### Design and Validation of Indel-Selective PCR assays for efficient genotyping of edits

Although mutagenesis by multiplex genome editing is effective, the spectrum of mutations generated can present challenges in screening for initial edits or downstream analyses to characterize functions of either individual or desired combinations of mutations. Large (>10-bp) deletion mutations can be easily screened by changes in PCR amplicon size, but small, often single-base indels that are the most common editing outcome are not detected by this approach. *In vitro* digestion assays with Cas9 ribonucleoproteins can reveal any type of mutation at a target site, but not the specific sequence changes, and are not amenable to high-throughput screening. Thus, we sought to design a general PCR-based approach to identifying and tracking small indel mutations.

To design the indel-selective assay, we leveraged the observation that single nucleotide insertions or deletions created by CRISPR-Cas9 can also disrupt base-pairing of PCR primers at adjacent nucleotides. We reasoned that three nucleotide mismatches at the 3’end of the primer would likely disable primer extension by DNA polymerase, even when the remainder of the primer sequence anneals. PCR primers can thus be designed where the anticipated indel mutation(s) and the two adjacent nucleotides will have three mismatches with the original genomic target but will perfectly match mutated DNA (Figure 4A & 4B). During PCR with a common reverse primer that matches both edited and reference alleles, the 3-nucleotide mismatch at the end of forward primers only allows proper primer annealing/extension to either the original or mutant sequence, therefore selectively amplifying one allele (Figure 4B, C). If the extension of the next two nucleotides on one DNA strand does not enable design of forward primers that are indel-selective, for example due to repeated nucleotides, the assay can be designed to operate in the reverse orientation with indel-selective reverse primers and a common forward primer. Notably, successful PCR amplification from the same DNA sample with both assays (original sequence & mutant) reveals heterozygous genotypes (Figure 4C).

**Figure 4:**
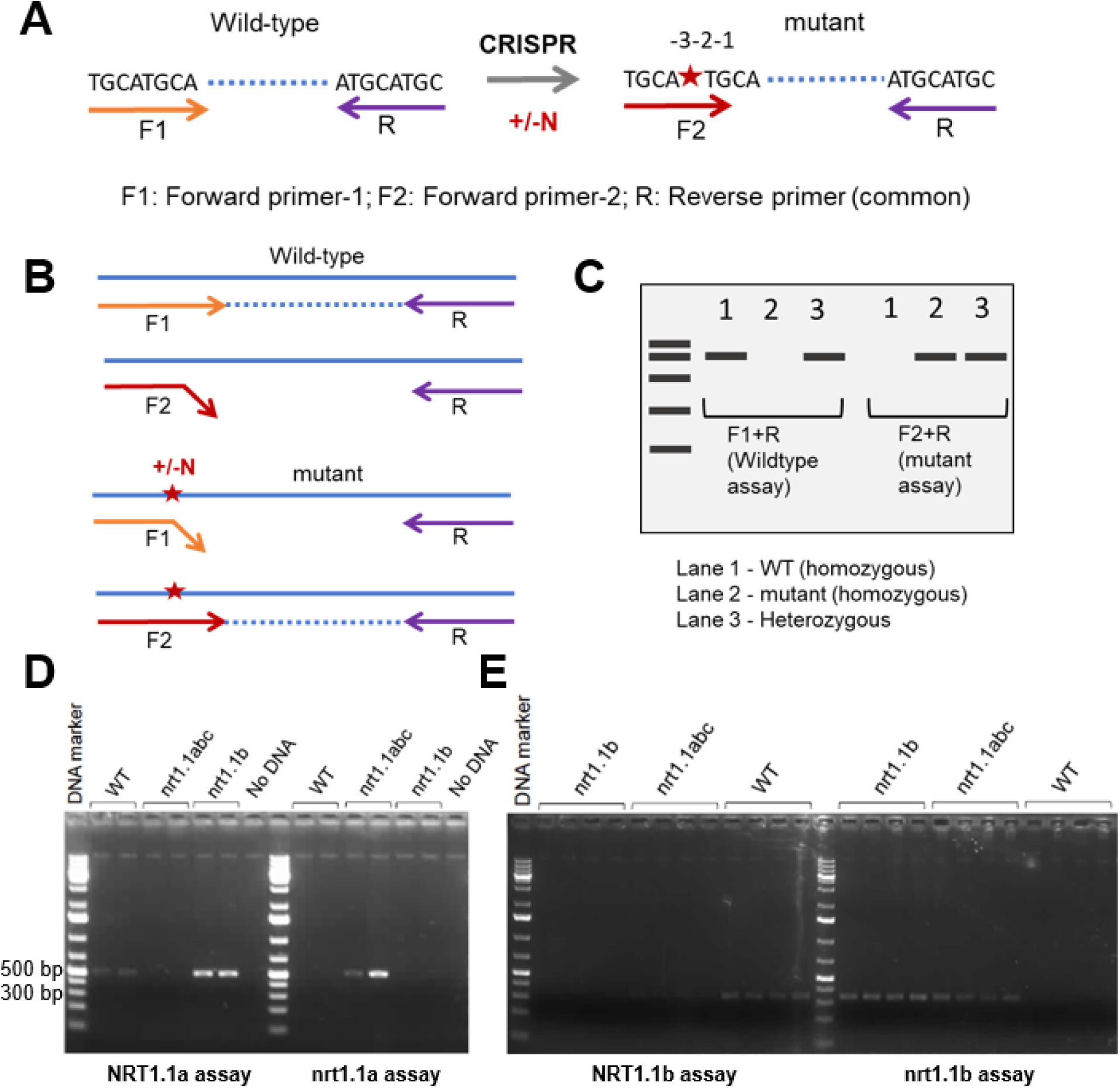
Indel-Selective PCR Assay. **(A)** Single base insertion/deletion (+/-N) created by CRISPR mutagenesis. To design wild type and indel allele specific forward primers (denoted by F1 & F2, respectively), single base indel is kept as third last nucleotide (shown as -3) from the 3’ end of the primers. Reverse primer is kept common in both assays. **(B)** Illustration of 3’ end of allele-specific primer annealing only to their respective DNA template resulting in PCR amplification. **(C)** Schematic of the agarose gel electrophoresis analysis for wild type and mutant assay revealing the zygosity and genotype of tested DNA samples. **(D)** Gel image displaying PCR amplification of desired size of DNA product for NRT1.1a and nrt1.1a genotyping assays. **(E),** Gel image displaying PCR amplification of desired size of DNA product for NRT1.1b and nrt1.1b genotyping assays.

We tested the indel-selective PCR approach to identify edited *nrt1.1* alleles. We used two events from the multiplex editing experiment, Event 233 that contains mutations in each of the three targeted *NRT1.1* genes (referred hereafter as *nrt1.1-abc*), and Event 1 that contains a single- base T insertion only in *NRT1.1B* (*nrt1.1b*). Assays using a forward primer matching the *NRT1.1a* reference allele amplified a DNA product of desired size from the wild-type control and *nrt1.1b* genomic DNA templates, but *nrt1.1abc* genomic DNA samples did not show any PCR amplification (Figure 4D). Conversely, assays with a forward primer matching the *nrt1.1a* mutation with a single base C insertion amplified a DNA product of expected size only with *nrt1.1abc* genomic DNA template. A similar assay was designed to selectively amplify either the *NRT1.1b* reference sequence or the *nrt1.1b* mutant alleles in both events, which share a T insertion at the same position. As expected, the assay for the *NRT1.1b* reference allele produced a ∼350 base pair DNA fragment only with wild type genomic DNA samples (Figure 4E). The *nrt1.1b* mutant assay amplified the expected size DNA fragments in both the *nrt1.1abc* and *nrt1.1b* genomic DNA samples but not with wild-type DNA (Figure 4E). Together, these observations demonstrate the indel-selective assays can identify edited mutations.

The above assays were subsequently employed to screen large progeny populations from events with multiplex edits, to identify individuals with single mutations, or desirable combinations of mutant alleles. During 2023 over both a summer field and fall greenhouse breeding cycles, 1152 plants were genotyped for *Nrt1.1a* and/or *Nrt1.1b* alleles (supplementary Figure S4). These plants derived from 46 families where the edited mutations are either being introgressed into additional genetic backgrounds or combined with other mutations in the common H99 background. Eight families were derived from crosses of the H99: *nrt1.1abc* triple mutant with H99, which subsequently segregated for individuals homozygous for either single *nrt1.1a* or tandemly-linked *nrt1.1b* and *nrt1.1c* mutations. From this genotyping campaign, we validated by Sanger sequencing five individuals identified as homozygous for either wild type or *nrt1.1* mutant alleles by our indel-selective PCR assays (supplementary Figure 5).

## Discussion

CRISPR/Cas9 genome editing technology has provided a significant boost towards crop improvement (Zhang *et al*., 2021; Zhu *et al*., 2020). Multiplexed genome editing using CRISPR/Cas9 has been achieved in maize but has been applied primarily to target two sites within the same gene. A number of studies have successfully mutagenized multiple genes (Char *et al*., 2017; Doll *et al*., 2019; Hurst *et al*., 2024; Liu *et al*., 2022, 2020; Lorenzo *et al*., 2023) but each have employed *Agrobacterium* transformation with constructs where gRNAs are expressed individually from RNA pol III promoters. The pioneering study by Qi *et al*. (2016) successfully generated edits in multiple genes by expressing gRNAs separated by tRNA spacers from the same transcript, which were subsequently processed by the endogenous tRNA biogenesis pathway. However, gRNA expression was driven by RNA pol III promoters and relatively large constructs were delivered by *Agrobacterium* to Hi-II embryogenic callus. We present here a ‘fast-edit’ approach for multiplexed-genome editing of maize inbred lines by biolistics delivery of a compact single DNA fragment containing all editing components to Type I embryogenic callus, combined with indel-selective PCR to efficiently identify edited mutations.

We demonstrate for the first time successful multiplex editing in maize using the streamlined vector design based on Csy4 processing (Čermák *et al*, 2017). The Csy4 multiplexing system offers two major advantages compared to RNA pol III promoters driving individual gRNAs. Firstly, monocot RNA pol III promoters such as U3 and U6 require a terminal 5’ G/A nucleotide in the gRNA for efficient expression, whereas the Csy4 multiplexing system bypasses this requirement and allows more flexibility in gRNA design and selection (Hsieh-Feng and Yang, 2020). Secondly, equal coexpression of gRNAs increases the probability each gRNA will be present in sufficient abundance to program Cas9 editing (Luo *et al*., 2021). Similar to prior multiplex editing experiments in maize, the majority of T0 events generated (9 of 13 for NRT1.1) harbored at least one edited mutation, and the majority of these mutations (10 of 19 for the NRT1.1 experiment, Table 3) were heritable. From the perspective of individual genes, editing frequency was on the order of 1 mutation per 100 initial embryos, or 1 heritable mutation from three to five transgenic events.

Although multiplex editing in maize has been achieved through biolistics delivery of editing components as separate fragments, this approach has so far been limited to targeting two sites within the same gene (Gao *et al*., 2020). Transformation of 12 different inbreds with four DNA fragments (Cas9, two gRNAs, *nptII* selectable marker gene) was enhanced by co-bombardment with two additional constructs (six fragments total) encoding morphogenic regulators, which may have reduced mutation transmission rates. In contrast, our methods use one DNA fragment of ∼12-kb that programmed efficient simultaneous editing of three paralagous *NRT1.1* genes, including recovery of a triple mutant line where all three genes were edited (*nrt1.1abc*). As has been observed in other maize editing experiments, many events are fully edited at the callus or T0 plant stage, indicating mutations were generated early in the transformation experiment. Although the lower frequency of quality single-copy events with biolistics delivery is certainly an issue in creating stable transgenic lines, a higher copy number of Cas9 and gRNA transgenes may benefit editing efficiency. Furthermore, the ability to deliver significantly more DNA molecules to single cells via biolistics compared to *Agrobacterium* may increase the potential for mutagenesis from transient expression of editing components.

After establishing a successful Csy4 multiplex genome editing pipeline for the H99 inbred known to be competent for transformation, we also succeeded in creating transgenic events and edited mutations for the ILP1 inbred line that has never been transformed previously. Transformation efficiencies for the *LW1* and *NRT1.1* editing experiments in H99 were in line with previous work at about 3% (Brettschneider *et al*., 1997). Transformation efficiency for the *LW1* editing experiment in ILP1 was slightly higher at about 7%, which we have subsequently observed can be further increased with media optimization (data not shown). The same methods have succeeded in inducing Type I embryogenic callus and fertile regenerated plants for a wide diversity of maize genotypes (Duncan et al., 1985; Hodges et al., 1986; Wan et al., 1995). The slower growth of Type I callus is a significant disadvantage with *Agrobacterium*-mediated transformation, because the rare transformed cells are less competitive in outgrowing *Agrobacterium* after co-cultivation, whereas this is not an issue with biolistics delivery. The methods described here open the possibility for direct editing of genotypes with other desirable traits, such as the unique kernel composition and nitrogen utilization phenotypes of ILP1 (Uribelarrea et al., 2007), thereby expanding the utility of CRISPR/Cas9 editing for maize functional genomics research.

The efficient mutagenesis of multiple targets achieved by multiplex editing complicates subsequent efforts to identify individual edits and track their inheritance. The *NRT1.1* gene editing experiment produced events with mutations in one gene, but the majority harbored mutations in more than one *NRT1.1* gene at a shared target sequence (e.g. *nrt1.1abc*), further complicating resolution of mutations in individual genes. As has been reported previously in plants (Bortesi *et al*., 2016), we observed single-base insertions as the most common type of edited mutation (Figure 4D). We demonstrate here that a single mismatched primer can selectively assay multiple classes of possible single-base indels. A few primers can be designed in advance to cover all possible anticipated single-base indels for a given target. Once the specific sequence change is determined, the indel-selective PCR assays can also be used for high-throughput genotyping of mutant populations.

The indel-selective PCR assay will enhance applications of genome editing to crop improvement, as it expands the capacity for simple low-cost PCR assays to detect single-base edited alleles with a clear path for commercialization in the US and other countries that have declared these mutations as exempt from regulatory oversight (Tachikawa and Matsuo, 2023). Depending on the country, methods using biolistics delivery without any extraneous or vector backbone DNA may also be favored over *Agrobacterium* for gaining eventual regulatory approvals. The genome-editing methods reported here are also not encumbered by intellectual property beyond patents active for CRISPR/Cas9 mutagenesis, so could be used by the global maize research community. Considering the entire pipeline from design and assembly of vectors, DNA delivery, transformation and recovery of events, and genotyping for mutant identification, our approach is easier to implement for scientists new to genome editing. Each of the experiments reported here were conducted primarily by graduate and undergraduate students, where edits were confirmed within six months of initial gRNA design and starting with only 500 immature embryos, and the subsequent tissue culture readily handled by a single scientist. The Csy4 multiplex vectors and indel-selective PCR assays can also be used for genome editing of related grasses. We have already demonstrated multiplex editing of the LW1 genes from polyploid *Miscanthus* (Trieu *et al*., 2022) and our group has recently applied this approach to generation and screening for single-base edits for sorghum genes using similar transformation methods (Brant *et al*., 2021). We anticipate the versatility of the methods described here will enhance the utility of genome editing for functional genomics research in maize and other grasses.

## Materials and Methods

### Guide RNA Design and CRISPR/Cas9 Expression Vector Construction

All guide RNAs (gRNAs) were designed using the CHOPCHOP web tool (https://chopchop.cbu.uib.no/). Default settings were used along with the B73 v4 genome for gRNA generation, efficiency scoring and off target prediction. All gRNAs chosen contained a GC content of 30-70%, a self-complementarity score of zero and no off-target sites with less than two mismatches. The off-target requirement was relaxed for *NRT1.1*−gRNA2 to allow for perfect matching at *NRT1.1A*, *NRT1.1B* and *NRT1.1C*. gRNAs designed from B73 sequence were confirmed to perfectly match target gene sequences present in the H99 and ILP1 genomes, as ascertained by perfect alignment of seedling RNASeq data from H99 (Hansey *et al*., 2012) or Illumina genome shotgun sequence from ILP1 (accession SRP229948 at NCBI Short Read Archive) with the B73 v4 reference assembly.

The constructs for multiplex CRISPR/Cas9 experiments were generated using the base plasmids and Golden-gate cloning system (Čermák *et al*., 2017) using appropriate primers. The specific plasmid components used were a monocot optimized Cas9 and Csy4 protein expressed under the maize Ubiquitin1 promoter and separated by a P2A self-cleaving peptide sequence (Addgene ID: 91036), a multiplexed gRNA cassette expressed with a Cestrum Yellow Leaf Curling Virus (CmYCLV) promoter and separated by Csy4 cleavage sites (Addgene ID: 91061), a blank homologous recombination module (Addgene ID: 91081) and a plasmid backbone containing the *NPTII* resistance gene driven by the 2X 35S promoter (Addgene ID: 91203).

### *In vitro* digestion of gene targets with Cas9 Ribonucleoproteins

*In vitro* Cas9 Ribonucleoprotein (RNP) digestion was performed as previously described with minor changes (Liang *et al*., 2017). Cas9-6X HIS protein was isolated following expression in *E. coli* using a Ni-NTA purification. Oligonucleotides for each designed gRNA were annealed, phosphorylated with T4 Polynucleotide Kinase (NEB Catalog #M0201S) and cloned into pT7- gRNA (Addgene Plasmid #46759). Synthetic RNA molecules of each gRNA target sequence and scaffold were created following NEB’s T7 RNA Synthesis Kit (Catalog #E2040S). Purified PCR products encompassing all gRNA target sites were amplified and eluted into RNase free water. The Cas9-gRNA digestion reaction was set up using Cas9 (1 µg), gRNA (375 ng), PCR product (200 ng), and 10X Cas9 reaction buffer (20mM HEPES, pH 7.5, 150 mM KC, 10mM MgCl2, 0.5mM DTT). The reaction was incubated at 37°C for 1 hour for Cas9 digestion and 25°C for 15 minutes after addition of proteinase K and at 56°C for 10 minutes. Digested products were run on a 2% agarose gel at 100V for 90 min and differences in band intensity were determined using ImageJ to calculate editing efficiencies.

### Plant materials, embryo excision and callus induction

All transformations in this study were conducted in maize inbred H99 and Illinois Low Protein (ILP1) lines, using the methods described in Shiva Prakash *et al*. (2008). Plants were either grown in a greenhouse or field, where ears from H99 were harvested between 10-12 days after pollination (DAP) and from ILP1 between 12-14 DAP. Prior to embryo excision, ears were surface sterilized by first drenching them in 70% ethanol, then submerging them into 30% bleach solution for 15 minutes, followed by multiple washes with sterile deionized water. After the final wash, the top 1-2mm of kernel crowns was carefully removed with a scalpel. Immature embryos ranging from 1.5 to 2.0 mm in length were then aseptically excised. The excised embryos were placed scutellar side up on N6E media supplemented with auxin, 3% sucrose, 100 mg/L myo-inositol, 2.27 g/L proline, 100 mg/L casein hydrolysate and 0.25% phytagel. For H99, the auxin source was 2,4 D (2 mg/L), whereas for ILP1 auxin was supplied as Dicamba (3.315mg/L) because of the higher frequency of regeneration with that media reported by Duncan *et al*. (1985). The plates were then incubated in the dark at 25°C.

### Maize transformation

Four-day old embryos or embryogenic calli were placed onto N6OSM medium four hours prior to bombardment. N6OSM media is N6 medium with the addition of 0.69 g/L of proline and concentrations of 3.64% sorbitol and 3.64% mannitol. The embryos/embryogenic calli were bombarded with the CRISPR/Cas9 vector of interest following precipitation of the plasmid onto 0.6 mM gold particles using spermidine and CaCl_2_ (Aulinger *et al*., 2003). After 16-24 hours of resting, the bombarded embryos/calli were transferred onto N6S medium containing 300 mg/L of paromomycin, plates were incubated in the dark at 25°C, and subcultured every two weeks until resistant calli could be identified.

After ∼8 weeks of selection, the resistant calli were moved onto Regeneration media-I (R-I) containing 300 mg/L of paromomycin and 5mg/L 6-Benzoaminopurine (6-BAP). After 3-4 days on this media, the calli were transferred onto R-I media containing 300 mg/L of paromomycin only and incubated in dark at 25°C. After 2-3 weeks the resistant calli were transferred into a growth chamber at 28°C with 16h/8h light/dark cycle. Calli pieces were initially exposed to a lower light intensity by covering them with three layers of paper towels, then acclimatized to higher light by removing one layer of paper towels every two to three days. Regenerating plantlets were transferred onto Regeneration media II (R-II) containing 150mg/L paramomycin in plastic cups (Solo). When plantlet shoots and roots grew to sufficient size in the plastic cups, they were transferred to moist soilless mix and allowed to develop under the same growth chamber conditions to acclimatize plants to lower relative humidity. Eventually, plants at the V3-V4 growth stage were transplanted to soil in 2L volume pots for T1 seed production in a greenhouse.

### Screening of putative transgenic lines

Genomic DNA of paromomycin-resistant calli or leaf tissue from regenerated plants was extracted and primers specific to Cas9 and guide RNA sequences (Supplementary Table 2) were used to identify events containing CRISPR/Cas9 components through PCR. Subsequently, an ELISA test for Cas9 (Epigentek Cat #: P-4060-96) was performed, according to the manufacturer’s protocol, to identify events expressing the protein. Sanger sequencing of PCR amplicons spanning guide RNA target sites were used to identify CRISPR/Cas9 induced mutations. Heterozygous, chimeric or complex alleles were analyzed using sequence trace decomposition as described in Brinkman *et al*. (Brinkman Eva Karina and van Steensel, 2019)

### Off-target CRISPR/Cas9 analysis

To identify and quantify any off-target mutations created during the *NRT1.1* CRISPR/Cas9 experiment, genomic DNA was extracted from *NRT1.1*-E1 callus tissue for deep sequencing. Genomic library preparation was conducted using the Nextera DNA Flex Library Prep Kit (Illumina Cat #: 20018704) according to the manufacturer’s protocol and sequencing was performed on an Illumina NovaSeq6000. Potential off target CRISPR/Cas9 recognition sites were initially predicted from DNA BLAST hits containing 3 or fewer mismatches with the NRT1.1 gRNAs within Oh43, the line most closely related to H99 for which a full genome sequence was available. Then, quality-filtered Illumina reads from the NRT1.1-E1 callus DNA were aligned to predicted off-targets, and alignments were manually inspected using Integrated Genomics Viewer for evidence of CRISPR/Cas9 mediated mutagenesis.

### Molecular characterization of edited lines and screening of progenies

For efficient genotyping of single-base indel edits, we designed Indel-Selective PCR assays. The sequences of the edited events obtained from Sanger sequencing of PCR amplicons were used to design allele-specific primers which enable specific PCR amplification of either edited or reference alleles. The indel-selective assay employs two forward primers, one matching the original genomic target sequence where the 3’nucleotide is positioned 2-bp away (-2 position) from the PAM, and a second similar forward primer where the third nucleotide from the 3’ end (- 5 position) matches the anticipated single-base indel. The two forward primers are combined in separate PCRs with a common reverse primer that perfectly matches both reference and mutant sequences located downstream of the target site, at a position that enables efficient amplification of a product that is easily visualized by agarose gel electrophoresis.

Seeds from crosses with edited mutations were sown either in potting soil and grown in the greenhouse, or in the field during the summer months. Four to six leaf samples were collected from each plant with a hand-held single hole paper punch into 96-well tubes, lyophyllized, and genomic DNA extracted with a modified CTAB method (Porebski *et al*., 1997). Genomic DNAs were assayed by PCR with indel-selective assays for both reference and mutant alleles, and PCR products visualized by agarose gel electrophoresis.

## Author Contribution

SPM, MNR, BR and SJ conceived the idea and designed the experiments. MNR, BR, SJ, performed the experiments and data analysis. ER helped with sequence analysis and off target analysis. SRC provided valuable advice and guidance on experimental design. PK helped with tissue culture. MNR, BR, SJ, PK and SPM wrote the manuscript. All authors read and approved the final manuscript.

## Acknowledgements

We thank Professor Dan Voytas for sharing the multiplexed Csy4-CRISPR/Cas9 plasmid used in the current work. We thank all the lab members of the Stephen Moose lab for insightful discussions and undergraduate students Camilla Macias and Jenna Donovan for their help during this work. We thank Sanger sequencing core of Roy J. Carver Biotechnology Center, University of Illinois at Urbana-Champaign for Sanger sequencing. This work was partially funded by the DOE Center for Advanced Bioenergy and Bioproducts Innovation (U.S. Department of Energy, Office of Science, Biological and Environmental Research Program under Award Number DE- SC0018420), the Denton E. and Betty Alexander Professorship in Maize Breeding and Genetics, and graduate fellowships to Brian Rhodes and Stephen Jinga from the Illinois Corn Marketing Board, and USDA-NIFA grant award number 2022-67013-37038. Any opinions, findings, and conclusions or recommendations expressed in this publication are those of the author(s) and do not necessarily reflect the views of either the U.S. Department of Energy or U.S. Department of Agriculture.

## Data availability

Sequence reads from edited H99 calli have been deposited at NCBI Short Read Archive under the accession number PRJNA1171432.

## References

Aesaert, S., Impens, L., Coussens, G., Van Lerberge, E., Vanderhaeghen, R., Desmet, L., et al. (2022) Optimized Transformation and Gene Editing of the B104 Public Maize Inbred by Improved Tissue Culture and Use of Morphogenic Regulators. Front Plant Sci, 13.

Aulinger, I.E., Peter, S.O., Schmid, J.E., and Stamp, P. (2003) Gametic embryos of maize as a target for biolistic transformation: comparison to immature zygotic embryos. Plant Cell Rep, 21, 585–591.

Bortesi, L., Zhu, C., Zischewski, J., Perez, L., Bassié, L., Nadi, R., et al. (2016) Patterns of CRISPR/Cas9 activity in plants, animals and microbes. Plant Biotechnol J, 14, 2203–2216.

Brant, E.J., Baloglu, M.C., Parikh, A., and Altpeter, F. (2021) CRISPR/Cas9 mediated targeted mutagenesis of LIGULELESS-1 in sorghum provides a rapidly scorable phenotype by altering leaf inclination angle. Biotechnol J, 16, 2100237.

Brettschneider, R., Becker, D., and Lörz, H. (1997) Efficient transformation of scutellar tissue of immature maize embryos. Theoretical and Applied Genetics, 94, 737–748.

Brinkman Eva Karina and van Steensel, B. (2019) Rapid Quantitative Evaluation of CRISPR Genome Editing by TIDE and TIDER. In: CRISPR Gene Editing: Methods and Protocols (Luo,Y., ed) , pp. 29–44. New York, NY: Springer New York.

Čermák, T., Curtin, S.J., Gil-Humanes, J., Čegan, R., Kono, T.J.Y., Konečná, E., et al. (2017) A Multipurpose Toolkit to Enable Advanced Genome Engineering in Plants. Plant Cell, 29, 1196–1217.

Char, S.N., Neelakandan, A.K., Nahampun, H., Frame, B., Main, M., Spalding, M.H., et al. (2017) An Agrobacterium-delivered CRISPR/Cas9 system for high-frequency targeted mutagenesis in maize. Plant Biotechnol J, 15, 257–268.

Cheng, C.-Y., Li, Y., Varala, K., Bubert, J., Huang, J., Kim, G.J., et al. (2021) Evolutionarily informed machine learning enhances the power of predictive gene-to-phenotype relationships. Nat Commun, 12, 5627.

Doll, N.M., Gilles, L.M., Gérentes, M.-F., Richard, C., Just, J., Fierlej, Y., et al. (2019) Single and multiple gene knockouts by CRISPR–Cas9 in maize. Plant Cell Rep, 38, 487–501.

Duncan, D.R., Williams, M.E., Zehr, B.E., and Widholm, J.M. (1985) The production of callus capable of plant regeneration from immature embryos of numerous Zea mays genotypes. Planta, 165, 322– 332.

Erenstein, O., Jaleta, M., Sonder, K., Mottaleb, K., and Prasanna, B.M. (2022) Global maize production, consumption and trade: trends and R&D implications. Food Secur, 14, 1295–1319.

Feng, C., Su, H., Bai, H., Wang, R., Liu, Y., Guo, X., et al. (2018) High-efficiency genome editing using a dmc1 promoter-controlled CRISPR/Cas9 system in maize. Plant Biotechnol J, 16, 1848–1857.

Feng, C., Yuan, J., Wang, R., Liu, Y., Birchler, J.A., and Han, F. (2016) Efficient Targeted Genome Modification in Maize Using CRISPR/Cas9 System. Journal of Genetics and Genomics, 43, 37–43.

Gao, H., Gadlage, M.J., Lafitte, H.R., Lenderts, B., Yang, M., Schroder, M., et al. (2020) Superior field performance of waxy corn engineered using CRISPR–Cas9. Nat Biotechnol, 38, 579–581.

Gordon-Kamm, W.J., Spencer, T.M., Mangano, M.L., Adams, T.R., Daines, R.J., Start, W.G., et al. (1990) Transformation of Maize Cells and Regeneration of Fertile Transgenic Plants. Plant Cell, 2, 603–618.

Hansey, C.N., Vaillancourt, B., Sekhon, R.S., de Leon, N., Kaeppler, S.M., and Buell, C.R. (2012) Maize (Zea mays L.) Genome Diversity as Revealed by RNA-Sequencing. PLoS One, 7, e33071-.

Hassan, M.M., Zhang, Y., Yuan, G., De, K., Chen, J.-G., Muchero, W., et al. (2021) Construct design for CRISPR/Cas-based genome editing in plants. Trends Plant Sci, 26, 1133–1152.

He, Y., Zhang, T., Yang, N., Xu, M., Yan, L., Wang, L., et al. (2017) Self-cleaving ribozymes enable the production of guide RNAs from unlimited choices of promoters for CRISPR/Cas9 mediated genome editing. Journal of Genetics and Genomics, 44, 469–472.

Hodges, T.K., Kamo, K.K., Imbrie, C.W., and Becwar, M.R. (1986) Genotype Specificity of Somatic Embryogenesis and Regeneration in Maize. Bio/Technology, 4, 219–223.

Hsieh-Feng, V. and Yang, Y. (2020) Efficient expression of multiple guide RNAs for CRISPR/Cas genome editing. aBIOTECH, 1, 123–134.

Hufford, M.B., Seetharam, A.S., Woodhouse, M.R., Chougule, K.M., Ou, S., Liu, J., et al. (2021) De novo assembly, annotation, and comparative analysis of 26 diverse maize genomes. Science (1979), 373, 655–662.

Hurst, J.P., Sato, S., Ferris, T., Yobi, A., Zhou, Y., Angelovici, R., et al. (2024) Editing the 19 kDa alpha-zein gene family generates non-opaque2-based quality protein maize. Plant Biotechnol J, 22, 946–959.

Ishida, Y., Saito, H., Ohta, S., Hiei, Y., Komari, T., and Kumashiro, T. (1996) High efficiency transformation of maize (Zea mays L.) mediated by Agrobacterium tumefaciens. Nat Biotechnol, 14, 745–750.

Kang, M., Lee, K., Finley, T., Chappell, H., Veena, V., and Wang, K. (2022) An Improved Agrobacterium- Mediated Transformation and Genome-Editing Method for Maize Inbred B104 Using a Ternary Vector System and Immature Embryos. Front Plant Sci, 13.

Liang, Z., Chen, K., Li, T., Zhang, Y., Wang, Y., Zhao, Q., et al. (2017) Efficient DNA-free genome editing of bread wheat using CRISPR/Cas9 ribonucleoprotein complexes. Nat Commun, 8, 14261.

Liu, H.-J., Jian, L., Xu, J., Zhang, Q., Zhang, Maolin, Jin, M., et al. (2020) High-Throughput CRISPR/Cas9 Mutagenesis Streamlines Trait Gene Identification in Maize[OPEN]. Plant Cell, 32, 1397–1413.

Liu, X., Zhang, S., Jiang, Y., Yan, T., Fang, C., Hou, Q., et al. (2022) Use of CRISPR/Cas9-Based Gene Editing to Simultaneously Mutate Multiple Homologous Genes Required for Pollen Development and Male Fertility in Maize. Cells, 11.

Lorenzo, C.D., Debray, K., Herwegh, D., Develtere, W., Impens, L., Schaumont, D., et al. (2023) BREEDIT: a multiplex genome editing strategy to improve complex quantitative traits in maize. Plant Cell, 35, 218–238.

Lowe, K., La Rota, M., Hoerster, G., Hastings, C., Wang, N., Chamberlin, M., et al. (2018) Rapid genotype “independent” Zea mays L. (maize) transformation via direct somatic embryogenesis. In Vitro Cellular & Developmental Biology - Plant, 54, 240–252.

Lowe, K., Wu, E., Wang, N., Hoerster, G., Hastings, C., Cho, M.-J., et al. (2016) Morphogenic Regulators Baby boom and Wuschel Improve Monocot Transformation. Plant Cell, 28, 1998–2015.

Luo, Y., Na, R., Nowak, J.S., Qiu, Y., Lu, Q.S., Yang, C., et al. (2021) Development of a Csy4-processed guide RNA delivery system with soybean-infecting virus ALSV for genome editing. BMC Plant Biol, 21, 419.

McCarty, N.S., Graham, A.E., Studená, L., and Ledesma-Amaro, R. (2020) Multiplexed CRISPR technologies for gene editing and transcriptional regulation. Nat Commun, 11, 1281.

Miao, J., Guo, D., Zhang, J., Huang, Q., Qin, G., Zhang, X., et al. (2013) Targeted mutagenesis in rice using CRISPR-Cas system. Cell Res, 23, 1233–1236.

Moose, S.P., Dudley, J.W., and Rocheford, T.R. (2004) Maize selection passes the century mark: a unique resource for 21st century genomics. Trends Plant Sci, 9, 358–364.

Plett, D., Toubia, J., Garnett, T., Tester, M., Kaiser, B.N., and Baumann, U. (2010) Dichotomy in the NRT Gene Families of Dicots and Grass Species. PLoS One, 5, e15289-.

Porebski, S., Bailey, L.G., and Baum, B.R. (1997) Modification of a CTAB DNA extraction protocol for plants containing high polysaccharide and polyphenol components. Plant Mol Biol Report, 15, 8–15.

Qi, W., Zhu, T., Tian, Z., Li, C., Zhang, W., and Song, R. (2016) High-efficiency CRISPR/Cas9 multiplex gene editing using the glycine tRNA-processing system-based strategy in maize. BMC Biotechnol, 16, 58.

Shiose, L., Moreira, J. dos R., Lira, B.S., Ponciano, G., Gómez-Ocampo, G., Wu, R.T.A., et al. (2024) A tomato B-box protein regulates plant development and fruit quality through the interaction with PIF4, HY5, and RIN transcription factors. J Exp Bot, 75, 3368–3387.

Shiva Prakash, N., Prasad, V., Chidambram, T.P., Cherian, S., Jayaprakash, T.L., Dasgupta, S., et al. (2008) Effect of promoter driving selectable marker on corn transformation. Transgenic Res, 17, 695–704.

Tachikawa, M. and Matsuo, M. (2023) Divergence and convergence in international regulatory policies regarding genome-edited food: How to find a middle ground. Front Plant Sci, 14.

Tang, X., Zheng, X., Qi, Y., Zhang, D., Cheng, Y., Tang, A., et al. (2016) A Single Transcript CRISPR-Cas9 System for Efficient Genome Editing in Plants. Mol Plant, 9, 1088–1091.

Trieu, A., Belaffif, M.B., Hirannaiah, P., Manjunatha, S., Wood, R., Bathula, Y., et al. (2022) Transformation and gene editing in the bioenergy grass Miscanthus. Biotechnology for Biofuels and Bioproducts, 15, 148.

Uribelarrea, M., Moose, S.P., and Below, F.E. (2007) Divergent selection for grain protein affects nitrogen use in maize hybrids. Field Crops Res, 100, 82–90.

Wan, Y., Widholm, J.M., and Lemaux, P.G. (1995) Type I callus as a bombardment target for generating fertile transgenic maize (Zea mays L.). Planta, 196, 7–14.

Wang, B., Lin, Z., Li, Xin, Zhao, Y., Zhao, B., Wu, G., et al. (2020) Genome-wide selection and genetic improvement during modern maize breeding. Nat Genet, 52, 565–571.

Wang, N., Ryan, L., Sardesai, N., Wu, E., Lenderts, B., Lowe, K., et al. (2023) Leaf transformation for efficient random integration and targeted genome modification in maize and sorghum. Nat Plants, 9, 255–270.

Wang, Z., Wan, L., Xin, Q., Zhang, X., Song, Y., Wang, P., et al. (2021) Optimizing glyphosate tolerance in rapeseed by CRISPR/Cas9-based geminiviral donor DNA replicon system with Csy4-based single-guide RNA processing. J Exp Bot, 72, 4796–4808.

Xing, H.-L., Dong, L., Wang, Z.-P., Zhang, H.-Y., Han, C.-Y., Liu, B., et al. (2014) A CRISPR/Cas9 toolkit for multiplex genome editing in plants. BMC Plant Biol, 14, 327.

Zhang, D., Zhang, Z., Unver, T., and Zhang, B. (2021) CRISPR/Cas: A powerful tool for gene function study and crop improvement. J Adv Res, 29, 207–221.

Zhang, J., Fengler, K.A., Van Hemert, J.L., Gupta, R., Mongar, N., Sun, J., et al. (2019) Identification and characterization of a novel stay-green QTL that increases yield in maize. Plant Biotechnol J, 17, 2272–2285.

Zhu, H., Li, C., and Gao, C. (2020) Applications of CRISPR–Cas in agriculture and plant biotechnology. Nat Rev Mol Cell Biol, 21, 661–677.

