## Supplementary figures and tables for "Efficient mutagenesis of maize inbreds using biolistics, multiplex CRISPR/Cas9 editing, and Indel-Selective PCR"

#### Supplementary Figure 1

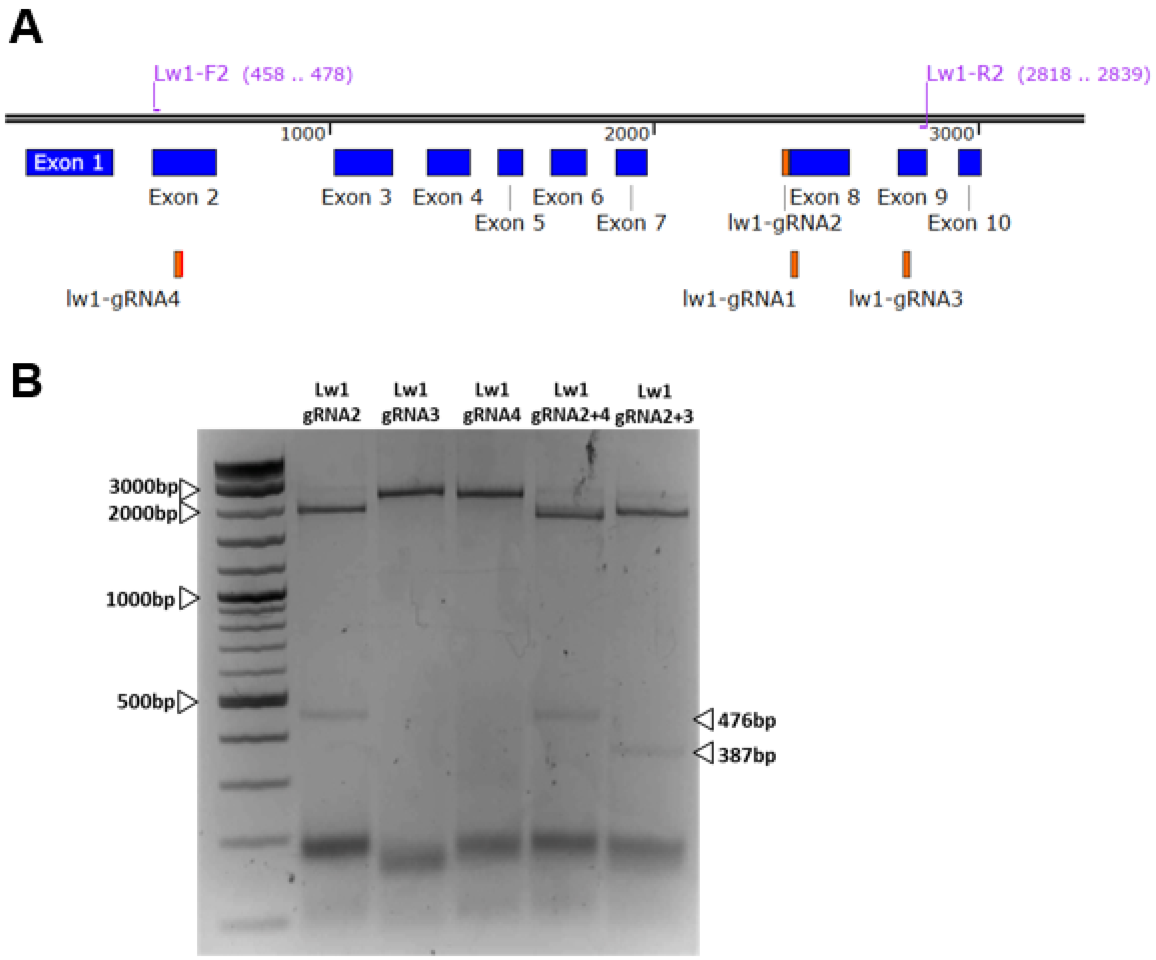


**Supplementary Figure 1: Demonstration of gRNA-targeted cleavage of LW1 DNA by Cas9 RNP in vitro digestion assays.** **(A)** Schematic of LW1 gene model with target sites for four gRNAs, and primers to amplify amplicons to test guide RNAs. **(B)** Gel electrophoresis of DNA fragments incubated with Cas9 RNPs loaded with either single or combinations of sgRNAs. Arrows indicate the sizes of fragments expected after cleavage. Guide RNAs 3 and 4 are close to primers to amplify amplicons and therefore did not produce detectable bands.

### Supplementary Figure 2

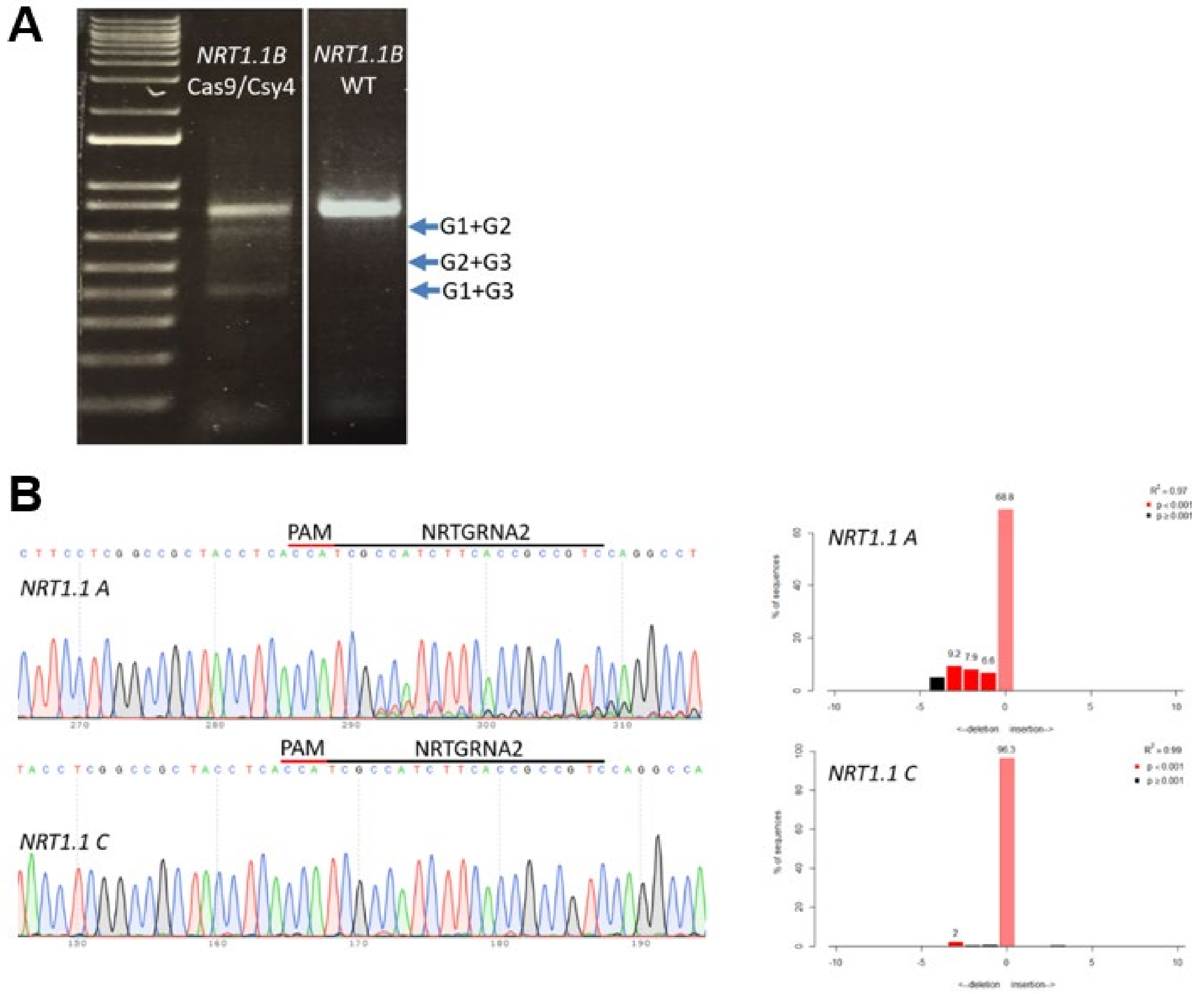


**Supplementary Figure 2:** Confirmation of NRT1.1-gRNA activity in maize leaf protoplasts. The NRT1.1 editing construct was transfected into protoplasts using PEG, genomic DNA extracted from protoplasts after 48 hours, followed by PCR amplification of DNA spanning the NRT1.1-gRNA target sites. (**A**) Gel electrophoresis shows DNA fragment sizes consistent with deletions in the *NRT1.1B* gene resulting from multiplex editing (blue arrows). (**B**) Sanger sequencing chromatograms and predicted proportion of editing for PCR products spanning the NRT1.1-gRNA2 cleavage site from the *NRT1.1A* and *NRT1.1C* genes.

### Supplementary Figure 3

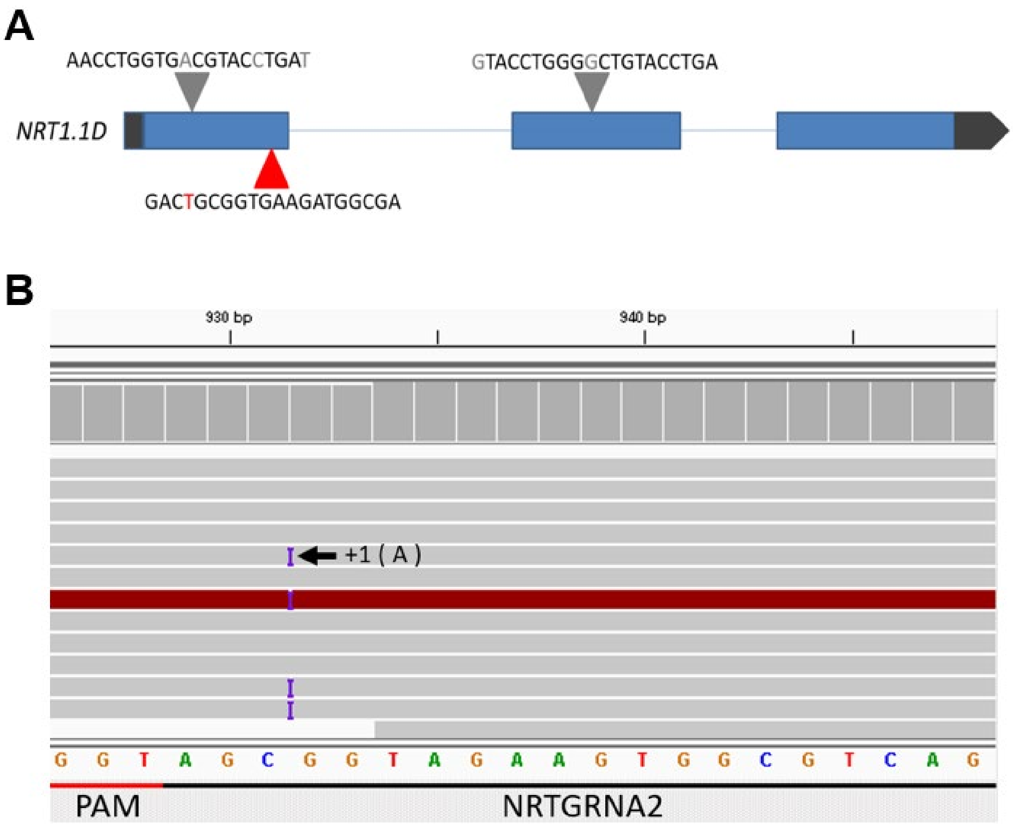


**Supplementary Figure 3:** Off target assessment of the CRISPR events. **(A)** Schematic representation of the off-target gene *NRT1.1D* gene model. The red triangle one base mismatch with the reference genome in the region complementary to guide RNA 2. (**B**) Integrated genome viewer (IGV) browser screenshot showing the insertion of nucleotide 'A' resulting in the off-target mutagenesis.

### Supplementary Figure 4

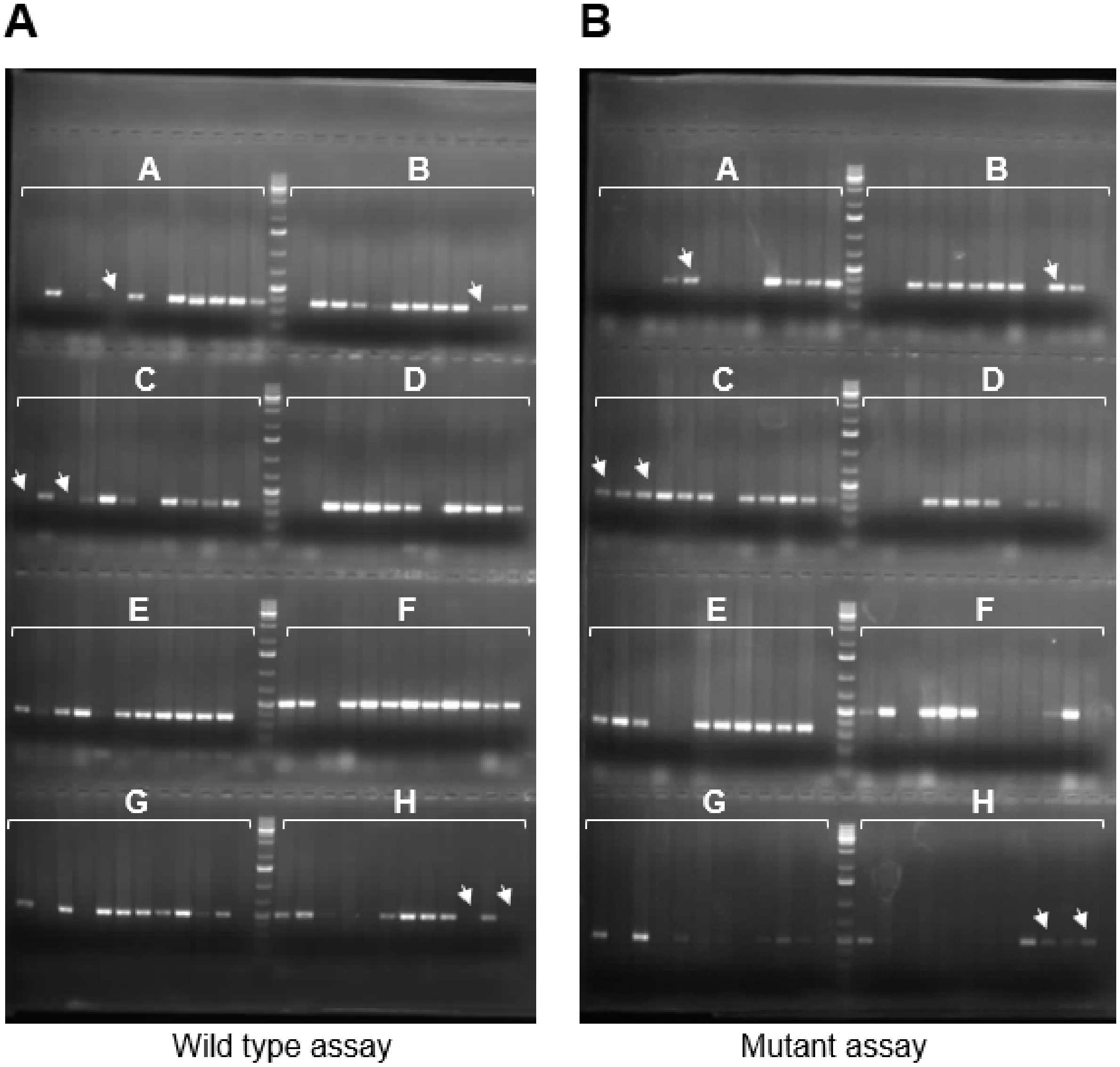


**Supplementary Figure 4: Representative gel picture of population segregating for *nrt1.1a* and *nrt1.1b*, to deconvolute homozygous mutant lines. (A)** Gel image depicting presence of *Nrt.1b* (rows A-E) and *Nrt1.1a* (rows F-H) wild type alleles in segregating maize lines. **(B)** Gel image displaying presence of nrt1.1b (rows A-E) and nrt1.1a (rows F-H) mutant alleles in segregating maize population. Arrow heads indicate the desired mutant lines showing bands for presence of mutant allele and absence of wild type allele.

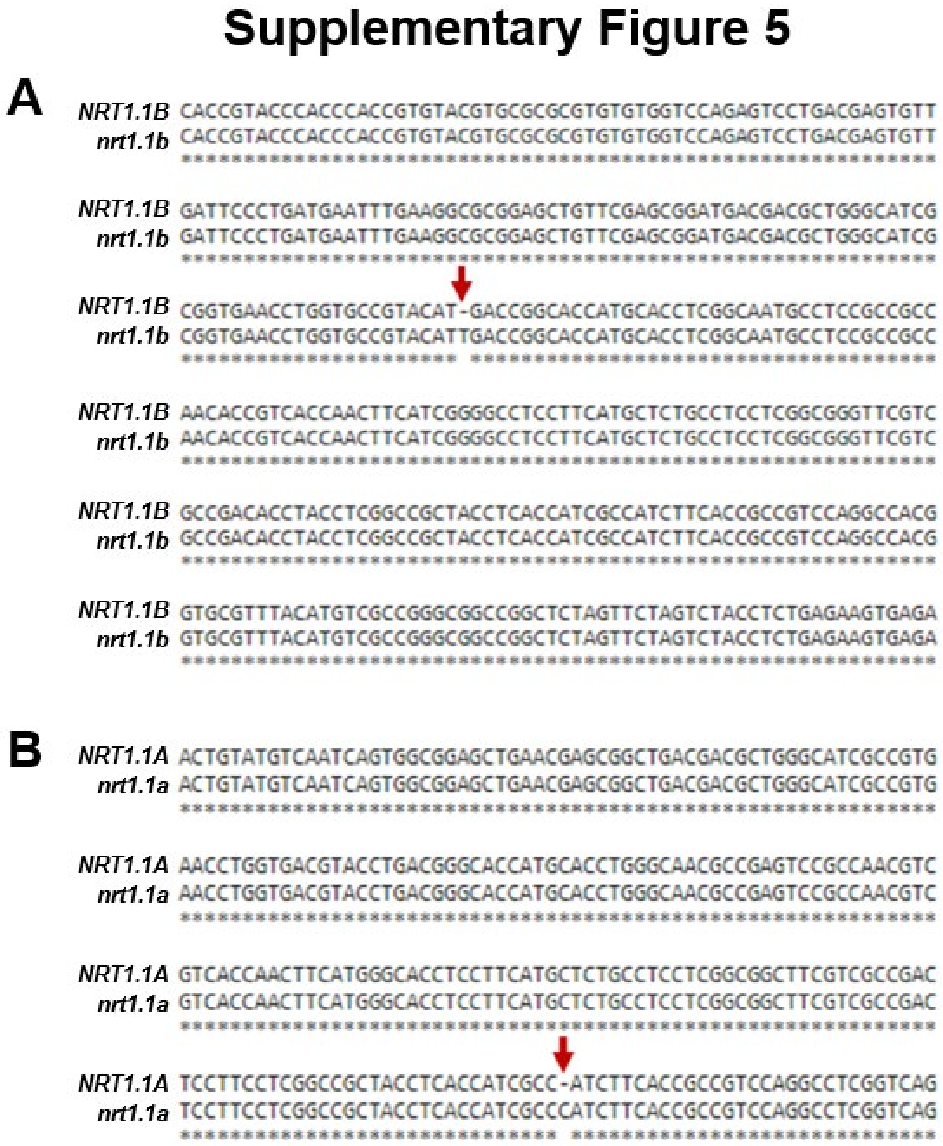


**Supplementary Figure 5: Validation of *NRT1.1B/nrt1.1b* and *NRT1.1A/nrt1.1a* IS-PCR genotyping results by Sanger sequencing. (A & B)** Sequence comparison of PCR amplified and purified amplicons from *NRT1.1* wild type and mutant plants identified by IASA genotyping method. Asterisks represent the common nucleotides and red arrows indicate single base insertion in *nrt1.1b* (A) and *nrt1.1a* mutant alleles (B).

**Supplementary Table 1: List of top 10 predicted off targets and results from sequencing for off-targets.**

| Off-target coordinates | Percent identity | Alignment length | Mismatches | Sequence | Edit | Notes |
|---|---|---|---|---|---|---|
| chr10:81131621-81131643 | 100 | 23 | 0 | CCATCGCCATCTTCACCGCCGTC | Y | *nrt1.1a* |
| chr1:95032630-95032652 | 100 | 23 | 0 | CCATCGCCATCTTCACCGCCGTC | Y | *nrt1.1b* |
| chr1:95089208-95089186 | 100 | 23 | 0 | CCATCGCCATCTTCACCGCCGTC | Y | *nrt1.1c* |
| chr1:1773343-1773365 | 95.65 | 23 | 1 | CCATCGCCATCTTCACCGCAGTC | Y | *nrt1.1d* |
| chr10:30075995-30076017 | 91.3 | 23 | 2 | CCATCGCCATCGTCACCGTCGTC | N | |
| chr8:61724083-61724061 | 91.3 | 23 | 2 | CCATCGCCATCTTCGCCGTCGTC | N | |
| chr6:36488051-36488073 | 91.3 | 23 | 2 | CCATCGCCATCGTCATCGCCGTC | N | |
| chr4:57413310-57413289 | 90.91 | 22 | 2 | CATCGCCATCTTCGCTGCCGTC | N | |
| chr9:14264778-14264799 | 90.91 | 22 | 2 | CATGGCCATCATCACCGCCGTC | N | |
| chr5:29734780-29734760 | 100 | 21 | 0 | CCATCGCCATCTTCACCGCCG | N | |

**Supplementary Table 2: List of primers used in this study.**

**For *LW1* editing experiment**

| Primer Name | Primer Sequence (5' --> 3') | Purpose |
|---|---|---|
| T7_lw1_g2_F | TAGGTGAGGAATCAAACATATGGT | gRNA cloning into T7 plasmid |
| T7_lw1_g2_R | AAACACCATATGTTTGATTCCTCA | |
| T7_lw1_g3_F | TAGGGAAAAATAACTGGTTACCCG | |
| T7_lw1_g3_R | AAACCGGGTAACCAGTTATTTTTC | |
| T7_lw1_g4_F | TAGGGGAGGCCTACGGGTTCTGCT | |
| T7_lw1_g4_R | AAACAGCAGAACCCGTAGGCCTCC | |
| CSY_LW1GRNA1 | TCGTCTCCATAGCATCTTGTCTGCCTATACGGCAGTGAAC | gRNA cloning into CRISPR/Cas9 expression vector cassette |
| REP_LW1GRNA1 | TCGTCTCACTATGTATCAGCGTTTTAGAGCTAGAAATAGC | |
| CSY_LW1GRNA2 | TCGTCTCCTTTGATTCCTCACTGCCTATACGGCAGTGAAC | |
| REP_LW1GRNA2 | TCGTCTCACAAACATATGGTGTTTTAGAGCTAGAAATAGC | |
| CSY_LW1GRNA3 | TCGTCTCCCAGTTATTTTTCCTGCCTATACGGCAGTGAAC | |
| REP_LW1GRNA3 | TCGTCTCAACTGGTTACCCGGTTTTAGAGCTAGAAATAGC | |
| CSY_LW1GRNA4 | TCGTCTCCCCGTAGGCCTCCCTGCCTATACGGCAGTGAAC | |
| REP_LW1GRNA4 | TCGTCTCAACGGGTTCTGCTGTTTTAGAGCTAGAAATAGC | |
| lw1-F1 | CACTTCATGGCCTTCAATAC | edit characterization |
| lw1-R2 | ACCTTATCTGGAGTTGAGGCAC | |
| lw1-F2 | GTCATCAAGACGCTCAAGGAG | |
| lw1-R5 | TGCAGTTAAGGCACGAACAC | |

**For *NRT1.1* editing experiment**

| Primer Name | Primer Sequence (5' --> 3') | Purpose |
|---|---|---|
| CSY_NRTGRNA1 | TCGTCTCCTTCACCGCCGTCCTGCCTATACGGCAGTGAAC | gRNA cloning into CRISPR/Cas9 expression vector cassette |
| REP_NRTGRNA1 | TCGTCTCATGAAGATGGCGAGTTTTAGAGCTAGAAATAGC | |
| CSY_NRTGRNA2 | TCGTCTCCCGGCACCAGGTTCTGCCTATACGGCAGTGAAC | |
| REP_NRTGRNA2 | TCGTCTCAGCCGTACATGACGTTTTAGAGCTAGAAATAGC | |
| CSY_NRTGRNA3 | TCGTCTCCAGACCCAGGTATCTGCCTATACGGCAGTGAAC | |
| REP_NRTGRNA3 | TCGTCTCAGTCTGTACCTGAGTTTTAGAGCTAGAAATAGC | |
| 086496_4F | GCCATGATCCTAGGTTGGTT | *NRT1.1A* edit characterization |
| 086496_4R | GTCGTCCCGAGTTTGTGG | |
| 161459_4F | GGATACTGCGTCGGATGG | *NRT1.1B* edit characterization |
| 161459_9F | TGTGTGGTCCAGAGTCCTGA | |
| 161459_5R | CGACACGCTGGACTTGAG | |
| 161459_9R | CCCATCGATCTCTTTGAACTAC | |

| | | |
|---|---|---|
| 112154_2F | TTTGTGGTCTCGTGAAGGTG | *NRT1.1C* edit characterization |
| 112154_5R | AATTAGCCATCAGCGTGTCC | |
| **For indel selective PCR genotyping** | | |
| **Primer name** | **Sequence** | **Purpose** |
| NRT1.1A_G2F | CCGCTACCTCACCATCGCCA | For *NRT1.1A* genotyping |
| nrt1.1a_e1_G2F | CCGCTACCTCACCATCGCCCA | |
| NRT1.1A_I2R | CCACCGCACCACCGTGCAT | |
| 161459 _4F | GGATACTGCGTCGGATGG | For *NRT1.1B* genotyping |
| NRT1.1b_3_G1R | GAGGTGCATGGTGCCGGTCATG | |
| nrt1.1b_3_G1R | GAGGTGCATGGTGCCGGTCAAT | |
